# Shifting microbial communities sustain multi-year iron reduction and methanogenesis in ferruginous sediment incubations

**DOI:** 10.1101/087783

**Authors:** Marcus S. Bray, Jieying Wu, Benjamin C. Reed, Cecilia B. Kretz, Keaton M. Belli, Rachel L. Simister, Cynthia Henny, Frank J. Stewart, Thomas J. DiChristina, Jay A. Brandes, David A. Fowle, Sean A. Crowe, Jennifer B. Glass

**Affiliations:** School of Biology, Georgia Institute of Technology, Atlanta, GA, 30332, USA; School of Earth and Atmospheric Sciences, Georgia Institute of Technology, Atlanta, GA, 30332, USA; Departments of Microbiology & Immunology and Earth, Ocean, & Atmospheric Sciences, University of British Columbia, Vancouver, BC, Canada; Research Center for Limnology, Indonesian Institute of Sciences, Cibinong, Indonesia; Skidaway Institute of Oceanography, Savannah, GA, 31411, USA; Department of Geology, University of Kansas, Lawrence, KS, 66045, USA

## Abstract

Reactive Fe(III) minerals can influence methane (CH_4_) emissions by inhibiting microbial methanogenesis or by stimulating anaerobic CH_4_ oxidation. The balance between Fe(III) reduction, methanogenesis, and methane oxidation in ferruginous Archean and Paleoproterozoic oceans would have controlled CH_4_ fluxes to the atmosphere, thereby regulating the capacity for CH_4_ to warm the early Earth under the Faint Young Sun. We studied CH_4_ and Fe cycling in anoxic incubations of ferruginous sediment from the ancient ocean analogue Lake Matano, Indonesia over three successive transfers (500 days total). Iron reduction, methanogenesis, methane oxidation, and microbial taxonomy were monitored in treatments amended with ferrihydrite or goethite. After three dilutions, Fe(III) reduction persisted only in bottles with ferrihydrite. Enhanced CH_4_ production was observed in the presence of goethite, highlighting the potential for reactive Fe(III)-oxides to inhibit methanogenesis. Supplementing the media with hydrogen, nickel and selenium did not stimulate methanogenesis. There was limited evidence for Fe(III)-dependent CH_4_ oxidation, although some incubations displayed CH_4_-stimulated Fe(III)-reduction. 16S rRNA profiles continuously changed over the course of enrichment, with ultimate dominance of unclassified members of the order Desulfuromonadales in all treatments. Microbial diversity decreased markedly over the course of incubation, with subtle differences between ferrihydrite and goethite amendments. These results suggest that Fe(III)-oxide mineralogy and availability of electron donors could have led to spatial separation of Fe(III)-reducing and methanogenic microbial communities in ferruginous marine sediments, potentially explaining the persistence of CH_4_ as a greenhouse gas throughout the first half of Earth history.

## INTRODUCTION

Elevated atmospheric methane (CH_4_; 100-1000 ppmv vs. ~2 ppmv in the modern atmosphere) likely played an important role in the first half of Earth history by helping warm Earth’s surface temperature enough to sustain liquid water under considerably lower solar radiation (Pavlov et al., 2000; Haqq-Misra et al., 2008; Kasting, 2005; Roberson et al., 2011). During this time, the main source of CH_4_ was likely hydrogenotrophic methanogenesis (CO_2_ + 4 H_2_ → CH_4_ + 2H_2_O; (Ueno et al., 2006; Battistuzzi et al., 2004) from anoxic oceans, which were ferruginous for most of the Archean and Paleoproterozoic eons (Poulton & Canfield, 2011). In these seas, a “ferrous wheel” would have cycled iron from dissolved Fe^2+^ to Fe(III) oxides via microbial photoferrotrophy (and/or abiotic photo-oxidation; Kappler et al., 2005; Crowe et al., 2008a), and then back to Fe^2+^ via microbial Fe(III) respiration (Craddock & Dauphas, 2011; Johnson et al., 2008; Vargas et al., 1998; Konhauser et al., 2005).

Ferruginous oceans could have influenced CH_4_ cycling by several mechanisms. It is well established that Fe(III)-reducing bacteria have higher affinity for H_2_ than hydrogenotrophic methanogens, and will therefore outcompete them in the presence of poorly crystalline Fe(III) oxides (e.g. ferrihydrite; Lovley & Phillips, 1987; Lovley & Goodwin, 1988; Zhou et al., 2014) [note that Fe(III)-reducing bacteria also outcompete acetoclastic methanogens (Lovley & Phillips, 1986), but acetoclastic methanogenesis likely evolved much later in Earth history (Fournier & Gogarten, 2008)]. In addition, evidence is accumulating that Fe(III) oxides can mediate or stimulate microbial CH_4_ oxidation, either as the direct oxidant (Ettwig et al., 2016; Gal’chenko, 2004), or indirectly by regenerating sulfate by oxidization of reduced sulfur compounds (Sivan et al., 2014).

The putative microbial metabolism of CH_4_ oxidation coupled to Fe(III) reduction is thermodynamically favorable with ferrihydrite (CH_4_ + 8 Fe(OH)_3_ + 15H^+^ → HCO_3_
^−^ + 8Fe^2+^ + 21H_2_O; ΔG_r_^0^ = -571 kJ mol^-1^ CH_4_) and goethite (CH_4_ + 8 FeOOH + 15H^+^ → HCO_3_^−^ + 8Fe^2+^ + 13H_2_O; ΔG_r_^0^ = -355 kJ mol^-1^ CH_4_) as terminal electron acceptors (Caldwell et al., 2008; Zehnder & Brock, 1980). Based on the chemical equations and free energy yields above, we would expect to observe a stoichiometric ratio of 1 CH_4_ oxidized per 8 Fe(III) reduced and preferential use of ferrihydrite over goethite as the electron acceptor. Accumulating geochemical evidence for microbial CH_4_ oxidation coupled to, or stimulated by, Fe(III) reduction is widespread across modern anoxic ecosystems and anaerobic digester communities (Sivan et al., 2011; Segarra et al., 2013; Beal et al., 2009; Amos et al., 2012; Riedinger et al., 2014; Noroi et al., 2013; Crowe et al., 2011; Sturm et al., 2015; Egger et al., 2015; Zehnder & Brock, 1980; Sivan et al., 2014; Fu et al., 2016; Rooze et al., 2016), and a recent study reported simultaneous CH_4_ oxidation and ferrihydrite reduction in a 1:8 ratio in an archaea-dominated enrichment culture (Ettwig et al., 2016).

Despite the possible importance of coupled Fe(III) and CH_4_ cycling in the Archean and Paleoproterozoic Eon, long-term studies of Fe(III) reduction under low organic carbon and high CH_4_ conditions remain sparse. Lake Matano, Indonesia is one of the only modern analogues for the ferruginous Archean ocean (Crowe et al., 2008a). Despite the abundance of Fe(III) oxides that might be expected to suppress methanogenesis, CH_4_ accumulates to 1.4 mM in anoxic deep waters (Crowe et al., 2008a; Crowe et al., 2011; Crowe et al., 2008b; Crowe et al., 2007a; Kuntz et al., 2015). Methanotrophy is a key carbon fixation process in Lake Matano’s oxic-anoxic transition zone, and the dearth of other oxidants (<100 nM nitrate and sulfate) suggests that Fe(III) might be the terminal electron acceptor in methanotrophy (Sturm et al., 2015; Crowe et al., 2011). In this study, we examined the influence of CH_4_ and Fe(III) mineral speciation on rates of Fe(III) reduction, methanogenesis, and CH_4_ oxidation, and microbial community composition, over three successive dilutions (500 total days of incubation) of anoxic Lake Matano sediments.

## MATERIALS AND METHODS

### Sample collection and storage

A 15-cm sediment core from 200 m water depth in Lake Matano, Sulawesi Island, Indonesia (2°26′S, 121°15′E; *in situ* sediment temperature ~27°C) was sampled in November 2010 and sub-sampled at 5 cm increments. Sediments from 0-5 and 5-10 cm depth were fluffy and black, and 10-15 cm was dark gray. Sediments were sealed in gas-tight bags with no headspace (Hansen et al., 2000) and stored at 4°C until incubations began in March 2015.

### Enrichment medium and substrate synthesis

A modified artificial freshwater medium lacking nitrate and sulfate was developed based on the pore water composition of Lake Matano sediments (S. A. Crowe and D. A. Fowle, unpublished work). The medium contained 825 µM MgCl_2_, 550 µM CaCO_3_, 3 mM NaHCO_3_, 3.5 µM K_2_HPO_4_, 5 µM Na_2_HPO_4_, 225 µM NH_4_Cl, 1 µM CuCl_2_, 1.5 µM Na_2_MoO_4_, 2.5 µM CoCl_2_, 23 µM MnCl_2_, 4 µM ZnCl_2_, 9.4 µM FeCl_3_ and 3 mM Na_2_NTA, 0.07 µM vitamin B_12_, 0.4 µM biotin, and 68.5 µM thiamine. Filter-sterilized vitamin solutions were added after autoclaving. Ferrihydrite (Fe(OH)_3_) and goethite (FeOOH) were synthesized as described in Schwertmann & Cornell (1991) and added to enrichments to 10 mM as described below.

### Inoculation of enrichment and amendments

The sediment was pre-treated for 36 days at 30°C in 100% N_2_ headspace to deplete endogenous organic carbon, electron donors, and reactive electron acceptors. After pre-treatment, sediment from the 0-5 cm depth layer was inoculated in a ratio of sediment to medium of 1:5 (v/v) in an anoxic chamber (97% N_2_ and 3% H_2_; Coy Laboratory Products, Grass Lake, MI, USA). Sediment slurry (35 mL) was aliquoted into 70 mL sterile serum bottles, stoppered with sterile butyl stoppers (Geo-Microbial Technologies, Ochelata, OK, USA; pre-boiled in 0.1 N NaOH), and crimped with aluminum seals. Ferric iron was added either as ferrihydrite or goethite to 10 mM. Bottles were purged with 99.99% N_2_ for 1 hr, and CH_4_ amendments were injected with 10 mL 99.99% CH_4_ and 5 mL 99% ^13^CH_4_ (Cambridge Isotope Laboratories, Tewksbury, MA, USA). Controls were autoclaved at 121°C for 1 hr on day 0 and again on day 6 of the 1° enrichment. All treatments were duplicated, and bottles were incubated in the dark at 30°C with shaking at 200 rpm.

After 50 days, the volume of all cultures was reduced to 5 mL, and 30 mL of fresh media was added to each bottle, constituting a 6-fold dilution. These 2**°** enrichments were amended with approximately 10 mM of either ferrihydrite or goethite. All bottles were purged with 99.99% N_2_ for 1 hr, and all bottles except N_2_ controls were injected with 8 mL 99.99% CH_4_ and 2 mL 99% ^13^CH_4_. Controls were autoclaved again at 121°C for 1 hr. DL-Methionine (10 µM) was added as a sulfur source. After 303 days, cultures were diluted 10-fold with fresh media into new serum bottles (3**°** enrichment) with the same substrate and headspace composition as the 2**°** enrichment. A schematic of the incubation and dilutions is shown at the top of Figures 1-3.

After an additional 220 days, goethite-amended N_2_ cultures were diluted 25-fold with fresh anoxic media into new serum bottles. Cultures received either 10 mM goethite or no Fe(III). A subset of cultures received 5 mL of 99.99% H_2_ (20% headspace) while all others had 100% N_2_ headspace. Controls were autoclaved at 121°C. After 48 days, an anoxic solution of nickel (Ni) and selenium (Se) was added to all bottles, yielding final concentrations of 1 µM Ni and 1 µM Se.

### HCl-extractable Fe^2+^ and Fe^3+^ and soluble Fe^2+^

Samples were taken from each bottle in the anoxic chamber using a 21-gauge needle (BD PrecisionGlide^TM^) and plastic syringe. Plasticware was stored in the anoxic chamber for at least 24 hr to minimize O_2_ sample contamination. For HCl-extractable Fe^2+^ analyses, 100 µL of sediment slurry was extracted with 400 µL 0.5 N HCl in the dark for 1 hr, followed by centrifugation at 10,000 x g for 1 min, injection of 10 µL of supernatant into 990 µL of 10 mM FerroZine reagent in 46 mM HEPES (pH 8.3), and measurement of absorbance at 562 nm (Stookey, 1970). For HCl-extractable Fe^3+^, 100 µL of sediment slurry was incubated overnight in 0.5 N HCl and 0.25 M NH_2_OH-HCl in the dark, followed by centrifugation and measurement as above, and subtraction of HCl-extractable Fe^2+^ as in Kostka & Luther (1994).

### Methane oxidation

Samples were collected for δ^13^C-DIC analysis by 0.2 µm membrane filtration of medium into crimp top autosampler vials (Thermo Scientific National Target LoVial) and analysis as described in Brandes (2009). Rates of ^13^CH_4_ oxidation to ^13^C-DIC were calculated over the linear period of δ^13^C-DIC increase based on the method in Scheller et al. (2016). First, the δ^13^C-DIC values were converted into fractional abundances (^13^F = (^13^C/^12^C+^13^C)), and then DIC production from CH_4_ oxidation was calculated using the following formula:

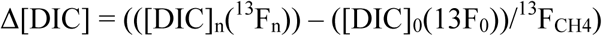

Where [DIC]_n_ and ^13^F_n_ are equal to the total DIC concentration (mM) and fractional abundance of ^13^C in the DIC at time n respectively. [DIC]_0_ and ^13^F_0_ are the total DIC concentration (mM) and fractional abundance of DIC at time 0 respectively, and ^13^F_CH4_ is the fractional abundance of ^13^C in the CH_4_.

### Headspace methane

Headspace (50 µL) was sampled using a gastight syringe and injected into a gas chromatograph (SRI Instruments 8610C, Torrance, CA, USA) with a HayeSep N column and flame ionization detector to measure headspace CH_4_ concentrations. A CH_4_ standard (1000 ppm, Airgas, USA) was used for calibration.

### Inductively coupled plasma mass spectrometry

Total dissolved Ni and Se concentrations were measured using inductively coupled plasma mass spectrometry (ICP-MS). In order to determine the amounts of Ni and Se supplied by the media and Fe(III) oxides, aliquots of media were dispensed in serum bottles, purged with 99.99% N_2_, and amended with 10 mM goethite or ferrihydrite in the same manner as enrichments. Stoppers were penetrated multiple times with 21 gauge stainless steel needles (BD PrecisionGlide^TM^) to mimic the effect of sampling on enrichment cultures. All samples for ICP-MS were filtered through 0.2 µm pore polypropylene syringe filters and diluted in 2% trace metal grade HNO_3_ (Fisher Scientific, Inc.) containing scandium and yttrium as internal standards to account for instrument drift. Calibration standards were prepared from certified stock solutions of Ni (CertiPREP) and Se (BDH), and a blank and calibration standard were measured periodically as quality controls. The measurement detection limits, calculated as 3 times the standard deviation of the blank (n=8), were 7 and 128 nM for Ni and Se, respectively.

### 16S rRNA gene amplicon sequencing

Samples (2 mL) of sediment used for inoculating incubations (hereafter, “sediment inoculum”) were taken in February 2015 (prior to pre-treatment) and after incubation for 15 days (1**°** enrichment), 72 days (2**°** enrichment) and 469 days (3**°** enrichment). Nucleic acid was extracted and purified using a MO BIO PowerSoil Isolation Kit following the manufacturer’s protocol and MO BIO UltraClean^®^ 15 Purification Kit. 16S rRNA gene amplicons were synthesized from extracted DNA with V4 region-specific barcoded primers F515 and R806 (Caporaso et al., 2011) appended with Illumina-specific adapters according to Kozich et al. (2013) using a Bio-Rad C1000 Touch Thermocycler and QIAGEN Taq PCR Master Mix. Thermal cycling conditions were as follows: initial denaturing at 94°C (5 min), 35 cycles of denaturing at 94°C (40 sec), primer annealing at 55°C (40 sec), and primer extension at 68°C (30 sec). Amplicons were checked for correct size (~400 bp) on a 1% agarose gel and purified using Diffinity RapidTips. Amplicon concentrations were determined on a Qubit^TM^ (ThermoFisher) fluorometer. Amplicons were pooled at equimolar concentrations (4 nmol), and sequenced on an Illumina MiSeq running MiSeq Control software v.2.4.0.4 using a 500 cycle MiSeq reagent kit v2 with a 5% PhiX genomic library control, as described by Kozich et al. (2013). Sequences were deposited as NCBI accession numbers SAMN04532568-04532641 and SAMN05915184-05915222.

### 16S rRNA gene amplicon sequence analysis

Demultiplexed amplicon read pairs were quality trimmed with Trim Galore (Babraham Bioinformatics) using a base Phred33 score threshold of Q25 and a minimum length cutoff of 100 bp. Reads were then analyzed using mothur (Schloss et al., 2009) following its MiSeq standard operating procedure. High quality paired reads were merged and screened to remove sequences of incorrect length and those with high numbers of ambiguous base calls. Merged reads were dereplicated and aligned to the ARB SILVA database (release 123; available at http://www.mothur.org/wiki/Silva_reference_alignment). Sequences with incorrect alignment and those with homopolymers longer than 8bp were filtered out. Unique sequences and their frequency in each sample were identified and then a pre-clustering algorithm was used to further de-noise sequences within each sample. Sequences were then chimera checked using UCHIME (Edgar et al., 2011). Reads were clustered into OTUs at 97% similarity based on uncorrected pairwise distance matrices. OTUs were classified using SILVA reference taxonomy database (release 123, available at http://www.mothur.org/wiki/Silva_reference_files). Chao 1 (species richness), phylogenetic diversity, and Shannon index (species evenness) estimates were generated using mothur after normalization to 4000 sequences per sample.

## RESULTS

### Iron reduction

*1° enrichment*. Over the first 10 days of incubation, HCl-extractable Fe^2+^ increased from 10 to 25 mM in ferrihydrite treatments (Fig. 1a) and from 10 to 20 mM in goethite treatments (1-2 mM d^-1^; Fig. 1b). From day 6 to 10, HCl-extractable Fe^3+^ (7 and 12 mM in ferrihydrite and goethite treatments, respectively) was completely consumed in all bottles except autoclaved controls with ferrihydrite (data not shown). Iron reduction rates were identical with and without CH_4_ (Fig. 1a, b). Initial autoclaving did not suppress Fe(III) reduction. A second round of autoclaving on day 6 slightly suppressed further activity. From day 10-28, HCl-extractable Fe^2+^ fluctuated in ferrihydrite treatments (Fig. 1a) and declined slightly in goethite treatments (Fig. 1b). Soluble Fe^2+^ was consistently <1% of HCl-extractable Fe^2+^, and sediment-free controls did not reduce Fe(III) (data not shown).

**Figure 1.**
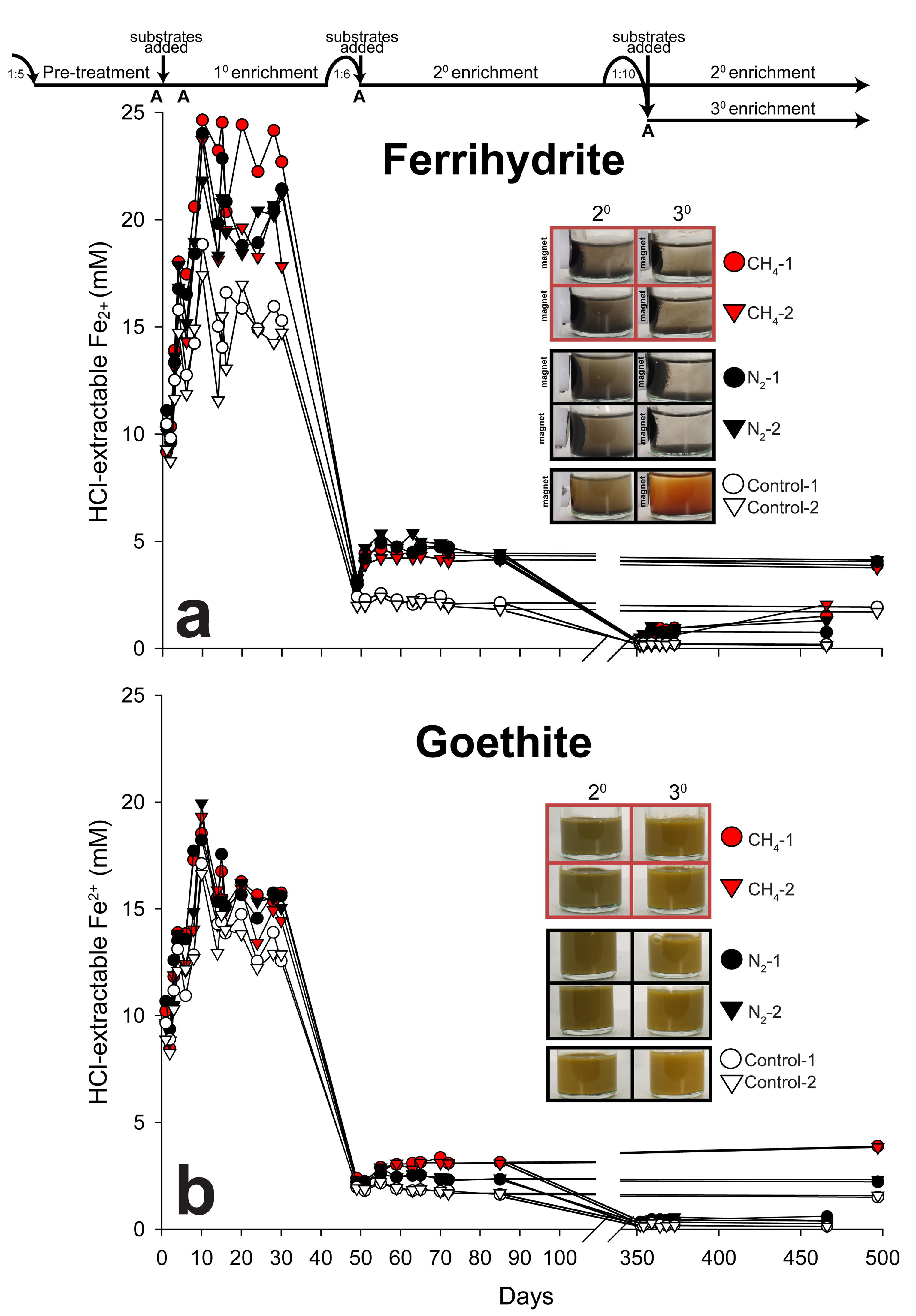
HCl-extractable Fe^2+^ for sediment enrichments with (a) ferrihydrite and (b) goethite over 497 days. Timeline at top shows transfer dates and dilution ratios. “A” represents days that controls were autoclaved. Red and black symbols represent treatments with and without CH_4_, respectively. White symbols represent autoclaved controls. All treatments were run in duplicate (circle and triangle symbols). Photos depict 2° and 3° enrichment bottles on day 497 with evidence for magnetic mineral formation in live treatments amended with ferrihydrite.

*2° enrichment*. After 1:6 dilution and 10 mM ferrihydrite addition on day 50, HCl-extractable Fe^2+^ increased from 3 to 4 mM over two days, and then remained constant through the final time point (day 497) in bottles with and without added CH_4_ (Fig. 1a). After 10 mM goethite addition on day 50, HCl-extractable Fe^2+^ increased from 2 to 3 mM after a two-day lag period. Thereafter, HCl-extractable Fe^2+^ rose to 4 mM by day 497 in goethite treatments with added CH_4_; without CH_4_, HCl-extractable Fe^2+^ dropped back to 2 mM (Fig. 1b). Autoclaved controls had no activity. Black magnetic minerals formed in all ferrihydrite treatments except autoclaved controls (Fig 1a). No magnetic minerals formed in goethite treatments (Fig. 1b).

*3° enrichment*. After 1:10 dilution and addition of 10 mM ferrihydrite on day 352, HCl-extractable Fe^2+^ doubled in the first week in N_2_ treatments and bottle CH_4_-1 (Fig. 1a). Bottle CH_4_-2 displayed similar activity after a two-week lag period. Over an additional 100 days (day 466), HCl-extractable Fe^2+^ increased to 2 mM. Goethite treatments and autoclaved controls had minimal activity (Fig. 1b). As in the 2*°* enrichment, magnetic minerals formed in the presence of ferrihydrite (Fig. 1a), but not goethite (Fig. 1b).

### Trace metal concentrations

Total dissolved Ni averaged 41 ± 20 nM in fresh basal growth media, and was neither affected by Fe(III) oxide additions nor by puncturing of stainless steel needles through stoppers into culture liquid (Table S1). Dissolution of ferrihydrite in HNO_3_ liberated significant Ni (2.5 μm; Table S1). Nickel was higher in enrichment cultures than in basal media: 96-286 and 54-134 nM with ferrihydrite and goethite, respectively; Table S2). Selenium was consistently below the detection limit (<128 nM) in growth media and enrichments culture.

### Methane production

Goethite treatments consistently displayed higher CH_4_ production than those with ferrihydrite (Fig. 2). In the 1° goethite enrichment, methanogenesis (13-19 µM CH_4_ d^-1^) coincided with the period of Fe(III) reduction, and stopped after HCl-extractable Fe^3+^ was completely consumed on day 10. Methanogenesis persisted throughout the 3° goethite enrichment (3 µM CH _4_ d^-1^). Negligible CH_4_ was produced in the presence of ferrihydrite, except for the final timepoint for bottle N_2_-2 in the 3° enrichment.

**Figure 2.**
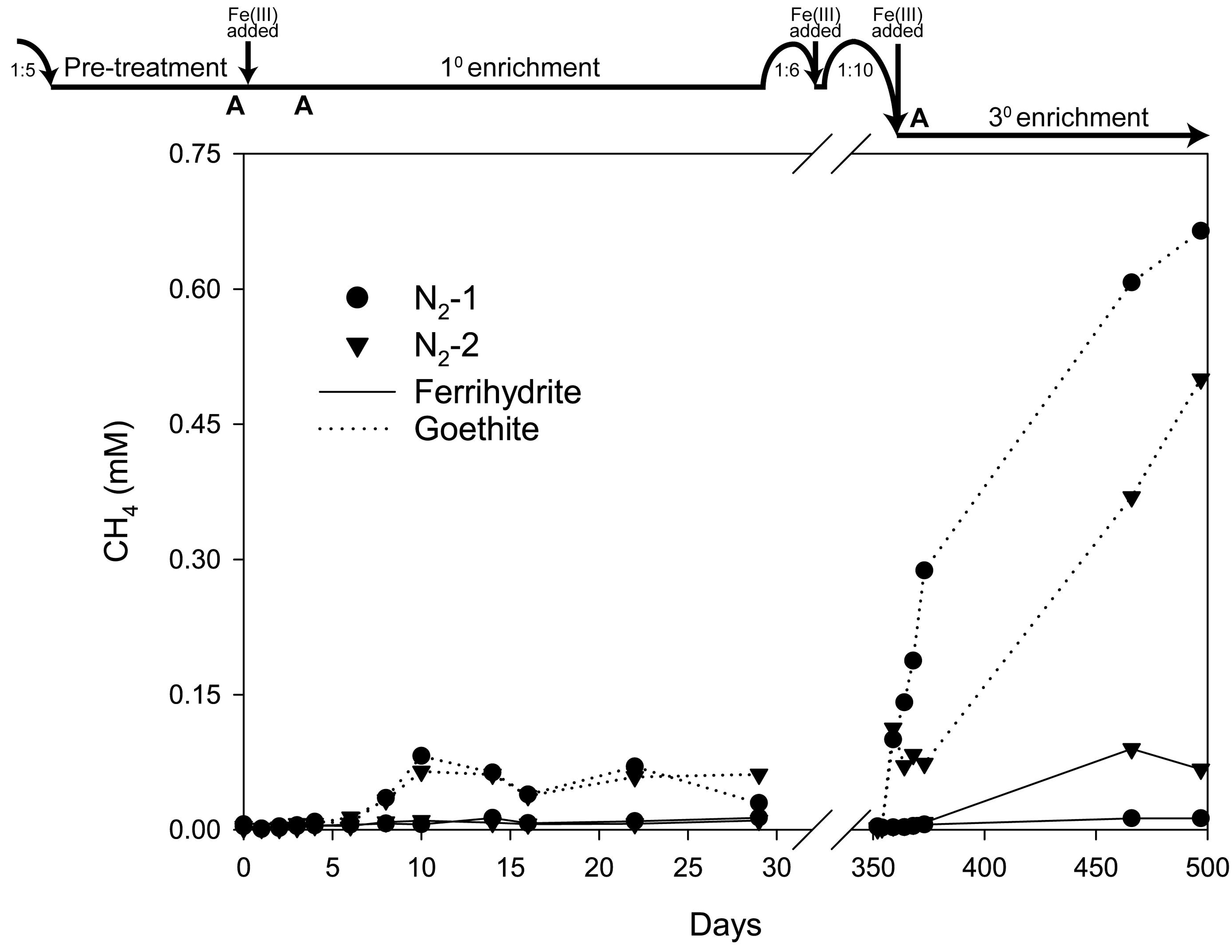
Accumulation of CH_4_ in the headspace of sediment enrichments. Timeline at top shows transfer dates and dilution ratios. Solid and dotted lines represent ferrihydrite and goethite treatments, respectively. All treatments were run in duplicate (circle and triangle symbols). Original headspace was 100% N_2_

In an additional 4° enrichment (day 571-663), we tested the effect of H_2_, Ni, and Se amendments, as well as no Fe(III) controls, on CH_4_ production in goethite treatments (Fig. S1). As in previous enrichments, Bottle 1 consistently produced more CH_4_ (17 µM CH_4_ d^-1^) than Bottle 2. There were no significant differences in CH_4_ production with and without 20% H_2_ headspace and 10 mM goethite. Like in previous enrichments with goethite, minimal Fe(III) reduction was observed (data not shown). No CH_4_ was produced in any of the treatments between day 619, when 1 µM Ni and Se were added, and day 663.

## Methane oxidation

*1° enrichment*. ^13^C incorporation into DIC began on day 6 in both ferrihydrite and goethite treatments and continued for the remainder of the sampling period (Fig. 3a, b). Ferrihydrite treatments showed lower ^13^C-DIC enrichment but higher total DIC production (totaling to 1-2 μM CH_4_ oxidized d^-1^) than goethite treatments, which had greatest δ^13^C enrichment but decreasing DIC concentrations, making calculation of CH_4_ oxidation rates impossible. Autoclaved controls showed neither ^13^C incorporation nor DIC production.

**Figure 3.**
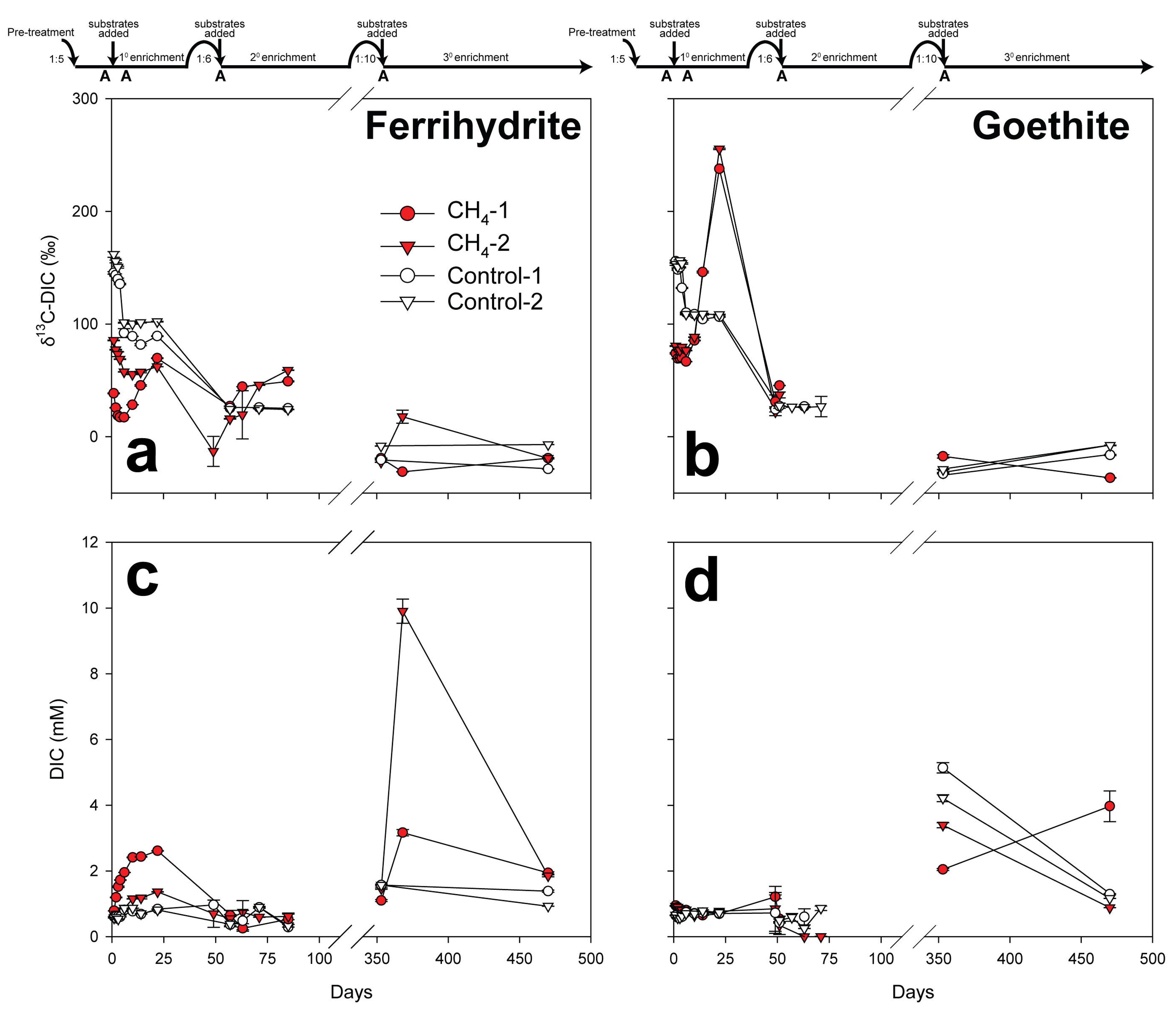
Dissolved inorganic carbon (DIC) isotopic composition and concentration for sediment enrichments amended with ^13^CH_4_ and either (a,c) ferrihydrite or (b,d) goethite. Timeline at top shows transfer dates and dilution ratios. “A” represents days that controls were autoclaved. Red and white symbols represent live treatments and autoclaved controls, respectively. Errors bars represent standard deviation of triplicate measurements. Calculated methane oxidation rates for the 1° enrichment were 1.7 and 1.1 μM CH_4_ d^-1^ ferrihydrite bottles 1 and 2, respectively and 0.2 and 0.8 μM CH_4_ d^-1^ for goethite bottles 1 and 2, respectively. Isotopic 694 data are not plotted for DIC concentrations 0.5 mM. Rate calculations were not possible for the 2° enrichment due to low/variable DIC.

*2° enrichment*. Both ferrihydrite treatments displayed ^13^C enrichment (Fig. 3a), but declining DIC, precluding calculation of CH_4_ oxidation rates. Initial pH of 8 declined to 7.6, 6.7 and 6 in the autoclaved, N_2_ and CH_4_ treatments, respectively. DIC in goethite treatments with CH_4_ dropped to undetectable values within three weeks, suggesting sampling or analytical error, which precluded accurate isotopic measurement at these time points. These data are thus not considered further. Autoclaved and N_2_ controls did not show pH changes.

*3° enrichment*. Bottle CH_4_-2 with ferrihydrite was the only treatment with significant ^13^C incorporation into DIC over the first 15 days (Fig. 3a). Over the same interval, DIC increased in both ferrihydrite-amended bottles (yielding CH_4_ oxidation rates of 32 and 7 µM d^-1^ in bottle 1 and 2, respectively) and pH dropped from 8.2 to 7.1 and 7.9 in bottles 1 and 2, respectively. By day 470, ^13^C enrichment and DIC concentrations in both ferrihydrite-amended bottles had returned to a level similar to that at the start of the 3° enrichment. Autoclaved controls did not exhibit any change in DIC and pH. Goethite treatments had initial DIC concentrations (3-5 mM) higher than those in previous enrichments. In the goethite-amended autoclaved controls and bottle CH_4_-1, DIC concentrations dropped over the 3° enrichment. Only goethite-amended bottle CH_4_-2 increased in DIC, without concurrent ^13^C enrichment. Large DIC variability implies that reported rates may be underestimates if declining pH led to outgassing of ^13^C-DIC into the headspace CO_2_ pool.

### Microbial taxonomy

*Inoculum*. 16S rRNA gene amplicons from the sediment inoculum were dominated by Bathyarchaeota (25%), formerly Miscellaneous Crenarchaeotal Group (MCG) and unclassified Archaea (11%; Fig. 4).

**Figure 4.**
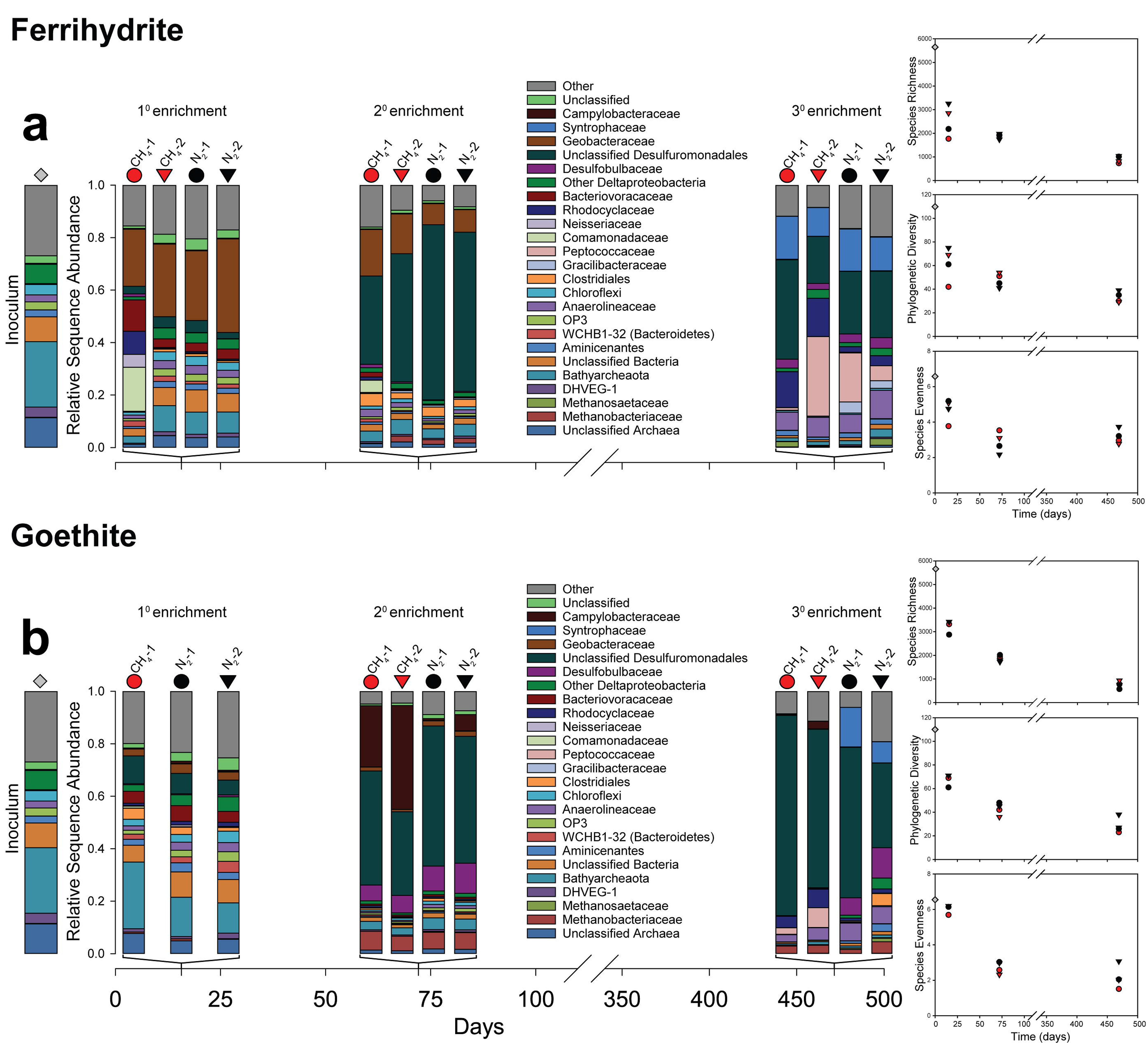
16S rRNA gene diversity and phylogenetic diversity for inoculum and sediment enrichments amended with (a) ferrihydrite and (b) goethite. Samples were taken on day 15 (1° enrichment), 72 (2° enrichment) and 469 (3° enrichment). Red and black symbols represent treatments with and without CH_4_, respectively. Gray diamonds represent inoculum samples. All treatments were run in duplicate (circle and triangle symbols). Species richness, phylogenetic diversity, and species evenness for the sediment inoculum and enrichments normalized to 4000 sequences per sample are shown to the right of bar charts.

*1° enrichment*. Species richness, evenness and phylogenetic diversity decreased relative to the inoculum in all treatments (Fig. 4). Geobacteraceae (Deltaproteobacteria) became dominant (22-36%) in all ferrihydrite treatments (Fig. 4a); the dominant OTU had 97% similarity to *Geothermobacter* sp. Ferrihydrite-amended bottle CH_4_-1 was enriched in the Betaproteobacteria, specifically Comamonaceadeae (17%) and Rhodocyclaceae (9%; Fig. 4a). Bathyarchaeota persisted in goethite treatments (11-25%; Fig. 4b).

*2° enrichment*. All treatments declined further in species richness, evenness, and phylogenetic diversity. Unclassified Desulfuromondales dominated both ferrihydrite and goethite enrichments (34-68%). The dominant OTU had 98% similarity to *Geobacter hephaestius/Geobacter lovleyi.* Geobacteraceae declined in ferrihydrite enrichments (2-18%; Fig. 4a). Campylobacteraceae (Epsilonproteobacteria), a trace constituent of the inoculum and 1° enrichment, were enriched in goethite treatments with CH_4_ (23-40%); the dominant OTU had 98% similarity to *Sulfurospirillum barnesii* (Fig. 4b). The most abundant methanogenic Euryarchaeota family, Methanobacteriaceae, comprised 1-2% and 6-7% of sequences in ferrihydrite and goethite treatments, respectively; the dominant OTU had 100% similarity to *Methanobacterium flexile*. Bathyarchaeota were depleted compared to the 1° enrichment (Fig. 4).

*3° enrichment*. Species richness, evenness, and phylogenetic diversity continued to decline (Fig. 4a, b). Unclassified Desulfuromondales dominated goethite treatments (32-76%; Fig. 4b), and were less abundant in ferrihydrite treatments (18-38%); as in the 2*°* enrichment, the dominant OTU had 98% identity to *Geobacter hephaestius/Geobacter lovleyi*. Rhodocyclaceae were more abundant in ferrihydrite treatments with CH_4_ (14-15%) than with N_2_ (2-4%; Fig 4a); the dominant OTU had 100% similarity to *Azospira oryzae/Dechlorosoma suillum*. Peptococcaceae (Firmicutes) were most abundant in bottle CH_4_-2 with ferrihydrite (30%); the dominant OTU had 96% similarity to uncultured members of the genus *Thermincola*. Syntrophaceae (Deltaproteobacteria) were enriched in all ferrihydrite treatments (11-16%) and goethite treatments with N_2_ (8-15%); the dominant OTU had 97% similarity to *Smithella propionica*. Methanobacteriaceae comprised 1-4% of sequences in goethite treatments and were absent from ferrihydrite treatments; as in the 2° enrichment, the dominant OTU had 100% identity to *Methanobacterium flexile*.

## DISCUSSION

### Fe(III) reduction rates in long-term ferruginous sediment incubations

Initial rates of HCl-extractable Fe^2+^ production (1-2 mM d^-1^) in the 1° enrichment were similar to those from freshwater wetlands with organic carbon as the electron donor (Roden & Wetzel, 2002; Jensen et al., 2003; Kostka et al., 2002). Despite replenishment of Fe(III) substrates, activity declined with each successive transfer, likely reflecting organic carbon limitation. The next most thermodynamically favorable electron donor, H_2_, could have been supplied by fermenters (such as Syntrophaceae in the 3° enrichment), but would ultimately still require a source of organic carbon. Some of our incubations display evidence for CH_4_, the next most thermodynamically favorable electron donor, as a source of electrons for Fe(III) reduction (e.g. higher Fe^2+^ yields with CH_4_ addition with ferrihydrite in 1° and 3° enrichments, and with goethite in the 2° enrichment; see further discussion below).

Higher Fe(III) reduction rates were maintained on ferrihydrite than goethite, consistent with its higher energetic yield and (typically) greater surface area. Magnetic mineral formation was likely due to adsorption of Fe^2+^ onto ferrihydrite followed by solid-state conversion of ferrihydrite to magnetite (Hansel et al., 2003). Since the HCl-extraction method does not dissolve magnetite and magnetite-adsorbed Fe^2+^ (Poulton & Canfield, 2005), it is possible that our Fe(III) reduction rates based on HCl-extractable Fe(II) production were underestimates of the total Fe(III) reduction.

### Fe(III) oxide mineralogy controls methane production and methanogen taxonomy

Our observation of higher rates of methanogenesis in goethite vs. ferrihydrite amendments is consistent with prior results showing that bacteria that reduce ferrihydrite better outcompete methanogenic archaea for H_2_ and acetate than those that reduce more crystalline Fe(III) oxides, including goethite (Lovley & Phillips, 1987; Lovley & Goodwin, 1988; Zhou et al., 2014; Hori et al., 2010; Roden & Wetzel, 1996). This out competition is also broadly supported by taxonomic shifts in our enrichment cultures. In particular, anaerobic heterotrophs such as *Geothermobacter sp.* (Kashefi et al., 2003) were enriched in ferrihydrite treatments by day 15 and may have outcompeted other microbes for organic carbon sources.

Higher abundances of Methanobacteriaceae (0.1-1% and 1-4% on days 15 and 469, respectively) in goethite than ferrihydrite treatments (≤0.1%) suggest that CH_4_ in goethite treatments came from the substrates used by Methanobacteriaceae (H_2_/CO_2_, formate, or CO). Addition of H_2_ did not stimulate additional methanogenesis in the 4° amendment, implying another limiting substrate or growth condition. The ferrihydrite treatment (bottle 2) that produced CH_4_ by day 469 contained 3% Methanosaetaceae; the most dominant OTU had 98% similarity to *Methanosaeta concilii,* in agreement with observations from the Lake Matano water column (Crowe et al., 2011). *Methanosaeta* spp. produce CH_4_ from acetate, or from H_2_/CO_2_ via direct interspecies electron transfer with *Geobacter* (Rotaru et al., 2014).

### Fe(III)-dependent CH_4_ oxidation

Enrichments were established under conditions thought to be favorable for Fe(III)-dependent CH_4_ oxidation, with Fe(III) oxides and CH_4_ as the most abundant electron acceptors and donors, respectively. In the 1° enrichment, incorporation of ^13^CH_4_ into DIC overlapped with the second phase of Fe(III) reduction (days 6-10), but calculating the stoichiometry of CH_4_ oxidized to Fe(III) reduced posed a challenge due to similar rates of Fe(III) reduction with and without added CH_4_ and in autoclaved controls. ^13^C-DIC enrichment from the back reaction of hydrogenotrophic methanogenesis (Zehnder & Brock, 1979) was ruled out because CH_4_ oxidation continued after Fe(III) reduction and methanogenesis stopped at day 10. Therefore, CH_4_ was likely oxidized by an electron acceptor other than Fe(III) (e.g. O_2_, Mn(IV), NO_x_^-^, SO_4_^2-^) in the 1° enrichment, likely supplied by residual sediment or inadvertent introduction of air.

During the first 15 days of the 3° enrichment, rates of CH_4_ oxidation and HCl-extractable Fe^2+^ production were similar (~10-20 µM d^-1^) and roughly consistent with the low rates presented in Ettwig et al. (2016) that yielded a 1:8 ratio. However, the lack of multiple time points for the interval of simultaneous Fe(III) reduction and CH_4_ oxidation, as well as similar initial rates of Fe(III) reduction with and without CH_4_ throughout this interval, prevent us from attributing this activity to Fe(III)-dependent CH_4_ oxidation with high confidence.

It is notable that the two incubations with the highest rates of CH_4_ oxidation (ferrihydrite bottle 1 in 1° and 3° enrichments) were also the only treatments with very different microbial community compositions relative to other bottles in the same enrichment. In the 1° enrichment, ferrihydrite bottle-1 was enriched in Betaproteobacteria (Comamonadaceae and Rhodocyclaceae). In the 3° enrichment, the CH_4_-1 sample had less Peptococcaceae than other ferrihydrite incubations. By day 469, the betaproteobacterium *Azospira oryzae/Dechlorosoma suillum*, a member of the Rhodocyclaceae family, was more abundant in both of the CH_4_ vs. N_2_ treatments. The potential role of this microbe in CH_4_ cycling remains unclear, as laboratory cultures of this species are not known to oxidize CH_4_. Notably, related members of the Betaproteobacteria, including the genera *Azospira* and *Comamonas* found here, are typically facultative anaerobes that can use alternative electron acceptors like NO_3_^-^, NO_2_^-^ or perchlorate (Willems, 2014; Reinhold-Hurek & Hurek, 2015). As such, these Betaproteobacteria are poised to respond to enhanced electron acceptor supply that accompanies pulse of O_2_. Anecdotally, Betaproteobacteria are frequently associated with environments that are characterized by fluctuating redox conditions and periodic exposure to O_2_ (Converse et al., 2015). Thus, their growth in our incubations may be a response to trace O_2_ introduction. It is also possible that the growth of novel organisms capable of high rates of Fe(III)-dependent CH_4_ oxidation was inhibited by other unidentified factors, potentially related to the batch-style incubations, the use of butyl rubber stoppers (Niemann et al., 2015), or the lack of a critical substrate in the enrichment medium.

### Effect of Fe(III) oxide and carbon substrates on microbial community diversity

The microbial community underwent multiple shifts over the 500-day incubation, with an overall decrease in species richness, evenness, and phylogenetic diversity, likely in response to declining organic carbon. By the 3° enrichment, species evenness was consistently lower in each goethite-amended treatment than in the respective ferrihydrite-amended treatment. This could mean that the greater energetic yield of ferrihydrite reduction fosters higher diversity, or that the higher reactivity of ferrihydrite allowed it to be utilized by more organisms than goethite.

All of the most enriched taxa in our enrichments comprised ≤0.1% of the inoculum community and have relatives that reduce Fe(III) in laboratory cultures. Within those taxa, the most abundant OTUs were closely related to organisms capable of Fe(III) reduction (*Geothermobacter sp.*, *Geobacter hephaestius/Geobacter lovleyi*, *Thermincola sp.*, and *Sulfurospirillum barnesii*) (Kashefi et al., 2003; Zavarzina et al., 2007; Stolz et al., 1999). Desulfuromonadales was the only metal-reducing taxon that was continuously present in all enrichments (3-11%, 34-53%, and 18-76% in the, 1°, 2°, and 3° enrichment respectively). Other taxa differed significantly in their abundance over the course of incubation. Geobacteraceae was enriched at day 15 with ferrihydrite (22-36%) but had declined in abundance by day 72 (8-18%). Still other taxa, including Rhodocyclaceae and Peptococcaceae, were enriched in the presence of ferrihydrite at day 469. In goethite treatments, Campylobacteraceae (23-40%) were enriched at day 72, but were minimal at day 469. The succession of different metal-reducing taxa may be due to the changing availability of electron donors (e.g. H_2_ and organic C). Enrichment of Syntrophaceae, known for their synthrophic fermentative interactions, suggests the establishment of syntrophy in the 3**°** enrichment in response to depletion of electron donors.

### Nickel sources

Enrichment cultures contained ~2-10x more total dissolved Ni than the basal growth medium. The inoculum (~60 nM in Lake Matano deep water; Crowe et al., 2008b) would not have significantly contributed to the Ni pool past the 1° enrichment, and repeated needle exposure had no effect on Ni concentrations. The Ni source to enrichment cultures was likely partial ferrihydrite dissolution, since ferrihydrite readily scavenges Ni from solution (Zegeye et al., 2012), while its dissolution liberates Ni (Table S1; Crowe et al., 2007b). Slow Ni leaching from silicate glass during extended contact between microbes and the serum bottles could have contributed another source of Ni in microbial enrichments vs. abiotic controls (Hausrath et al., 2007).

### Geobiological implications

Our results point to a mineralogical control on Fe(III) reduction, methanogenesis, and microbial community composition and diversity, under conditions of severe organic carbon limitation. These conditions likely existed in Archean and Paleoproterozoic oceans with relatively low amounts of primary production (Knoll et al., 2016; Farquhar et al., 2011). We posit that the relative abundance and distribution of Fe(III) phases in marine sediments would have impacted methanogenesis rates in the Archean and Paleoproterozoic. Sediments below shallow water columns were likely fed by abundant amorphous Fe(III) from photoferrotrophic activity, resulting in rapid sedimentation of amorphous Fe(III) phases (e.g. ferrihydrite). These Fe(III) oxides could have supported diverse Fe(III)-reducing communities that outcompeted other taxa such as methanogens for limited carbon and nutrients.

Conversely, slow deposition and aging of ferrihydrite to goethite could have limited both the abundance and diversity of Fe(III)-reducing microbes in sediments, allowing for more organic carbon remineralization via methanogenesis than Fe(III) reduction, as recently calculated for Lake Matano (Kuntz et al., 2015; Crowe et al., 2011). In the open ocean, organic carbon and Fe(III) would likely have been consumed before reaching sediments, leaving behind more crystalline Fe(III) phases. Importantly, the role of Fe(III)-driven CH_4_ oxidation appears limited given our experimental results, although we cannot rule out this pathway given that some of our data suggest it may operate at low rates.

Availability of trace metal nutrients is another important consideration in potential controls on ancient CH_4_ and Fe cycling. Measurements of Ni/Fe ratios in ancient marine sediments indicate that total dissolved Ni decreased from ~400 nM before 2.7 Ga to 200 nM between 2.7-2.5 Ga, to modern levels of 2-11 nM at ~0.5 Ga, assuming that the Fe(III) minerals in Archean sediments were of biological origin (Konhauser et al., 2009; Eickhoff et al., 2014; Konhauser et al., 2015). It is likely that abundant Ni would have been bound and sequestered in Fe(III)-oxide rich sediments. Rapid and widespread Archean redox cycling of Fe(III) could have served as constant source of Ni for methanogenic communities. The influence of changing availability of Se (Stüeken et al., 2015) and other trace nutrients on methanogenesis rates through time remains open for further exploration.

Overall, our results support a model for a sustained CH_4_ greenhouse in the Archean and Paleoproterozoic due to emissions from ferruginous oceans with spatially segregated habitats of bacterial reduction of reactive Fe(III) oxides and methanogenesis in the presence of less reactive Fe(III) phases. Rates of Fe(III) deposition, aging, and recrystallization may thus have played an important role in regulating the preservation of sedimentary Fe(III), the production of CH_4_, and the ecology and diversity of the biosphere during the first half of Earth history. By the mid-Proterozoic, rising seawater sulfate likely stimulated anaerobic CH_4_ oxidation, thereby minimizing marine CH_4_ emissions and the CH_4_ greenhouse (Olson et al., 2016).

Figure S1. **Accumulation of CH_4_ in the headspace during the 4° enrichment (days 571-663)**. All treatments were run in duplicate (circle and triangle symbols). The arrow represents addition of 1 μM Ni and Se on day 619. Original headspace was 100% N_2_ (black symbols) or 80 N_2_/20% H_2_ (blue symbols). White symbols represent autoclaved controls. Solid and dashed lines represent goethite and no Fe(III) treatments, respectively.

**Table S1.** **Total dissolved Ni (nM) in basal media with and without Fe(III) oxides**. Samples were prepared in the same manner as enrichment cultures. A subset of samples was acidified in HNO_3_^−^ prior to filtering and measurement. Error is reported as the standard deviation of triplicate measurements. “n.d.” indicates not detectable.

| Number of needle exposures | Medium | Medium +10 mM ferrihydrite | Medium +10 mM goethite | Medium +10 mM ferrihydrite + HNO<sub>3</sub> | Medium +10 mM goethite + HNO<sub>3</sub> |
| --- | --- | --- | --- | --- | --- |
| 1 | 41 ± 20 | 32 ± 1 | 50 ± 36 | 2365 | 65* |
| 2 | 44 ± 13 | 33 ± 11 | 52 ± 31 | n.d. | n.d. |
| 4 | 39 ± 9 | 30 ± 17 | 33 ± 10 | n.d. | n.d. |
| 6 | 41 ± 18 | 22 ± 19 | 33 ± 15 | 2385 | 605 |
| 8 | 41 ± 12 | 40 ± 19 | 42 ± 23 | n.d. | n.d. |
| 10 | 46 ± 16 | 38 ± 27 | 35 ± 4 | n.d. | n.d. |

\*Counts for this measurement were below detection limit of the instrument.

**Table S2.** **Total dissolved Ni(nM) in 2° and 3° enrichment cultures.**

| Treatment |  | Enrichment |  |
| --- | --- | --- | --- |
|  |  | 2° | 3° |
| Ferrihydrite | CH<sub>4</sub>-1 | 130 | 241 |
|  | CH<sub>4</sub>-2 | 150 | 118 |
|  | N<sub>2</sub>-1 | 202 | 96 |
|  | N<sub>2</sub>-2 | 286 | 254 |
| Goethite | CH<sub>4</sub>-1 | 104 | 101 |
|  | CH<sub>4</sub>-2 | 56 | 117 |
|  | N<sub>2</sub>-1 | 53 | 134 |
|  | N<sub>2</sub>-2 | 54 | 90 |

## Acknowledgements

This research was funded by NASA Exobiology grant NNX14AJ87G. Support was also provided by a Center for Dark Energy Biosphere Investigations (NSF-CDEBI OCE-0939564) small research grant, and supported by the NASA Astrobiology Institute (NNA15BB03A). SAC was supported through NSERC CRC, CFI, and Discovery grants. We thank Miles Mobley and Johnny Striepen for assistance with laboratory incubations, and Nadia Szeinbaum, Liz Percak-Dennett, Martial Taillefert, and Joel Kostka for helpful discussions.

## References

Amos R, Bekins B, Cozzarelli I, Voytek M, Kirshtein J, Jones E, Blowes D (2012) Evidence for iron-mediated anaerobic methane oxidation in a crude oil-contaminated aquifer. Geobiology, 10, 506–517.

Battistuzzi FU, Feijao A, Hedges SB (2004) A genomic timescale of prokaryote evolution: insights into the origin of methanogenesis, phototrophy, and the colonization of land. BMC Evolutionary Biology, 4, 44.

Beal EJ, House CH, Orphan VJ (2009) Manganese– and iron-dependent marine methane oxidation. Science, 325, 184–187.

Brandes JA (2009) Rapid and precise δ13C measurement of dissolved inorganic carbon in natural waters using liquid chromatography coupled to an isotope-ratio mass spectrometer. Limnology and Oceanography: Methods, 7, 730–739.

Caldwell SL, Laidler JR, Brewer EA, Eberly JO, Sandborgh SC, Colwell FS (2008) Anaerobic oxidation of methane: mechanisms, bioenergetics, and the ecology of associated microorganisms. Environmental Science & Technology, 42, 6791–6799.

Caporaso JG, Lauber CL, Walters WA, Berg-Lyons D, Lozupone CA, Turnbaugh PJ, Fierer N, Knight R (2011) Global patterns of 16S rRNA diversity at a depth of millions of sequences per sample. Proceedings of the National Academy of Sciences, 108, 4516–4522.

Converse BJ, Mckinley JP, Resch CT, Roden EE (2015) Microbial mineral colonization across a subsurface redox transition zone Frontiers in microbiology 6.

Craddock PR, Dauphas N (2011) Iron and carbon isotope evidence for microbial iron respiration throughout the Archean. Earth and Planetary Science Letters, 303, 121–132.

Crowe S, Katsev S, Leslie K, Sturm A, Magen C, Nomosatryo S, Pack M, Kessler J, Reeburgh W, Roberts J (2011) The methane cycle in ferruginous Lake Matano. Geobiology, 9, 61–78.

Crowe S, Roberts J, Weisener C, Fowle D (2007a) Alteration of iron-rich lacustrine sediments by dissimilatory iron-reducing bacteria. Geobiology, 5, 63–73.

Crowe SA, Jones C, Katsev S, Magen CD, O'neill AH, Sturm A, Canfield DE, Haffner GD, Mucci A, Sundby BR (2008a) Photoferrotrophs thrive in an Archean Ocean analogue. Proceedings of the National Academy of Sciences, 105, 15938–15943.

Crowe SA, O'neill AH, Katsev S, Hehanussa P, Haffner GD, Sundby B, Mucci A, Fowle DA (2008b) The biogeochemistry of tropical lakes: A case study from Lake Matano, Indonesia. Limnology Oceangraphy, 53, 319–331.

Crowe SA, O'neill AH, Kulczycki E, Weisener CG, Roberts JA, Fowle DA (2007b) Reductive dissolution of trace metals from sediments. Geomicrobiology Journal, 24, 157–165.

Edgar RC, Haas BJ, Clemente JC, Quince C, Knight R (2011) UCHIME improves sensitivity and speed of chimera detection. Bioinformatics, 27, 2194–2200.

Egger M, Rasigraf O, Sapart CLJ, Jilbert T, Jetten MS, RöCkmann T, Veen Van Der C, BâNdă N, Kartal B, Ettwig KF (2015) Iron-mediated anaerobic oxidation of methane in brackish coastal sediments. Environmental Science & Technology, 49, 277–283.

Eickhoff M, Obst M, Schröder C, Hitchcock AP, Tyliszczak T, Martinez RE, Robbins LJ, Konhauser KO, Kappler A (2014) Nickel partitioning in biogenic and abiogenic ferrihydrite: the influence of silica and implications for ancient environments. Geochim. Cosmochim. Acta, 140, 65–79.

Ettwig KF, Zhu B, Speth D, Keltjens JT, Jetten MSM, Kartal B (2016) Archaea catalyze iron-dependent anaerobic oxidation of methane. Proceedings of the National Academy of Sciences, 113, 12792–12796.

Farquhar J, Zerkle AL, Bekker A (2011) Geological constraints on the origin of oxygenic photosynthesis. Photosynthesis Research, 107, 11–36.

Fournier GP, Gogarten JP (2008) Evolution of acetoclastic methanogenesis in Methanosarcina via horizontal gene transfer from cellulolytic Clostridia. Journal of Bacteriology, 190, 1124–1127.

Fu L, Li S-W, Ding Z-W, Ding J, Lu Y-Z, Zeng RJ (2016) Iron reduction in the DAMO/Shewanella oneidensis MR-1 coculture system and the fate of Fe(II). Water Research, 88, 808–815.

Gal'chenko V (2004) On the problem of anaerobic methane oxidation. Microbiology, 73, 599–608.

Hansel CM, Benner SG, Neiss J, Dohnalkova A, Kukkadapu RK, Fendorf S (2003) Secondary mineralization pathways induced by dissimilatory iron reduction of ferrihydrite under advective flow. Geochimica et Cosmochimica Acta, 67, 2977–2992.

Hansen JW, Thamdrup B, Jørgensen BB (2000) Anoxic incubation of sediment in gas-tight plastic bags: a method for biogeochemical process studies. Marine Ecology Progress Series, 208, 273–282.

Haqq-Misra JD, Domagal-Goldman SD, Kasting PJ, Kasting JF (2008) A revised, hazy methane greenhouse for the Archean Earth. Astrobiology, 8, 1127–1137.

Hausrath EM, Liermann LJ, House CH, Ferry JG, Brantley SL (2007) The effect of methanogen growth on mineral substrates: will Ni markers of methanogen based communities be detectable in the rock record? Geobiology, 5, 49–61.

Hori T, Müller A, Igarashi Y, Conrad R, Friedrich MW (2010) Identification of iron-reducing microorganisms in anoxic rice paddy soil by 13C-acetate probing. The ISME journal, 4, 267–278.

Jensen MM, Thamdrup B, Rysgaard S, Holmer M, Fossing H (2003) Rates and regulation of microbial iron reduction in sediments of the Baltic-North Sea transition. Biogeochemistry, 65, 295–317.

Johnson CM, Beard BL, Roden EE (2008) The iron isotope fingerprints of redox and biogeochemical cycling in modern and ancient Earth. Annual Reviews of Earth and Planetary Sciences, 36, 457–493.

Kappler A, Pasquero C, Konhauser KO, Newman DK (2005) Deposition of banded iron formations by anoxygenic phototrophic Fe(II)-oxidizing bacteria. Geology, 33, 865–868.

Kashefi K, Holmes DE, Baross JA, Lovley DR (2003) Thermophily in the Geobacteraceae: Geothermobacter ehrlichii gen. nov., sp. nov., a Novel Thermophilic Member of the Geobacteraceae from the “Bag City” Hydrothermal Vent. Applied and Environmental Microbiology, 69, 2985–2993.

Kasting JF (2005) Methane and climate during the Precambrian era. Precambrian Research137, 119–129.

Knoll AH, Bergmann KD, Strauss JV (2016) Life: the first two billion years. Phil. Trans. R. Soc. B, 371, 20150493.

Konhauser K, Newman D, Kappler A (2005) The potential significance of microbial Fe(III) reduction during deposition of Precambrian banded iron formations. Geobiology, 3, 167–177.

Konhauser KO, Pecoits E, Lalonde SV, Papineau D, Nisbet EG, Barley ME, Arndt NT, Zahnle K, Kamber BS (2009) Oceanic nickel depletion and a methanogen famine before the Great Oxidation Event. Nature, 458, 750–753.

Konhauser KO, Robbins LJ, Pecoits E, Peacock C, Kappler A, Lalonde SV (2015) The Archean nickel famine revisited. Astrobiology, 15, 804–815.

Kostka JE, Luther GW (1994) Partitioning and speciation of solid phase iron in saltmarsh sediments. Geochemica Cosmochimica Acta, 58, 1701–1710.

Kostka JE, Roychoudhury A, Van Cappellen P (2002) Rates and controls of anaerobic microbial respiration across spatial and temporal gradients in saltmarsh sediments. Biogeochemistry, 60, 49–76.

Kozich JJ, Westcott SL, Baxter NT, Highlander SK, Schloss PD (2013) Development of a dual-index sequencing strategy and curation pipeline for analyzing amplicon sequence data on the MiSeq Illumina sequencing platform. Applied and Environmental Microbiology, 79, 5112–5120.

Kuntz LB, Laakso TA, Schrag DP, Crowe SA (2015) Modeling the carbon cycle in Lake Matano. Geobiology, 13, 454–461.

Lovley D, Phillips E (1986) Organic matter mineralization with reduction of ferric iron in anaerobic sediments. Applied and Environmental Microbiology, 51, 683–689.

Lovley DR, Goodwin S (1988) Hydrogen concentrations as an indicator of the predominant terminal electron-accepting reactions in aquatic sediments. Geochimica et Cosmochimica Acta, 52, 2993–3003.

Lovley DR, Phillips EJ (1987) Competitive mechanisms for inhibition of sulfate reduction and methane production in the zone of ferric iron reduction in sediments. Applied and Environmental Microbiology, 53, 2636–2641.

Niemann H, Steinle L, Blees J, Bussmann I, Treude T, Krause S, Elvert M, Lehmann MF (2015) Toxic effects of lab-grade butyl rubber stoppers on aerobic methane oxidation. Limnology and Oceanography: Methods, 13, 40–52.

Noroi KA, Thamdrup B, Schubert CJ (2013) Anaerobic oxidation of methane in an iron-rich Danish freshwater lake sediment. Limnology and Oceanography, 58, 546–554.

Olson SL, Reinhard CT, Lyons TW (2016) Limited role for methane in the mid-Proterozoic greenhouse. PNAS, 113, 11447–11452.

Pavlov AA, Kasting JF, Brown LL, Rages KA, Freedman R (2000) Greenhouse warming by CH4 in the atmosphere of early Earth. Journal of Geophysical Research: Planets, 105, 11981–11990.

Poulton SW, Canfield DE (2005) Development of a sequential extraction procedure for iron: implications for iron partitioning in continentally derived particulates. Chemical Geology, 214, 209–221.

Poulton SW, Canfield DE (2011) Ferruginous conditions: a dominant feature of the ocean through Earth's history. Elements, 7, 107–112.

Reinhold-Hurek B, Hurek T (2015) Azospira. Bergey's Manual of Systematics of Archaea and Bacteria, 1–3.

Riedinger N, Formolo M, Lyons T, Henkel S, Beck A, Kasten S (2014) An inorganic geochemical argument for coupled anaerobic oxidation of methane and iron reduction in marine sediments. Geobiology, 12, 172–181.

Roberson AL, Roadt J, Halevy I, Kasting J (2011) Greenhouse warming by nitrous oxide and methane in the Proterozoic Eon. Geobiology, 9, 313–320.

Roden EE, Wetzel RG (1996) Organic carbon oxidation and suppression of methane production by microbial Fe (III) oxide reduction in vegetated and unvegetated freshwater wetland sediments. Limnology and Oceanography, 41, 1733–1748.

Roden EE, Wetzel RG (2002) Kinetics of microbial Fe (III) oxide reduction in freshwater wetland sediments. Limnology and Oceanography, 47, 198–211.

Rooze J, Egger M, Tsandev I, Slomp CP (2016) Iron-dependent anaerobic oxidation of methane in coastal surface sediments: Potential controls and impact. Limnology and Oceanography.

Rotaru A-E, Shrestha PM, Liu F, Shrestha M, Shrestha D, Embree M, Zengler K, Wardman C, Nevina KP, Lovley DR (2014) A new model for electron flow during anaerobic digestion: direct interspecies electron transfer to Methanosaeta for the reduction of carbon dioxide to methane. Energy & Environmental Science, 7, 408–415.

Scheller S, Yi H, Chadwick GL, Mcglynn SE, Orphan VJ (2016) Artificial electron acceptors decouple archaeal methane oxidation from sulfate reduction. Science, 351, 703–707.

Schloss PD, Westcott SL, Ryabin T, Hall JR, Hartmann M, Hollister EB, Lesniewski RA, Oakley BB, Parks DH, Robinson CJ (2009) Introducing mothur: open-source, platform-independent, community-supported software for describing and comparing microbial communities. Applied and environmental microbiology, 75, 7537–7541.

Segarra KEA, Comerford C, Slaughter J, Joye SB (2013) Impact of electron acceptor availability on the anaerobic oxidation of methane in coastal freshwater and brackish wetland sediments. Geochimica et Cosmochimica Acta, 115, 15–30.

Sivan O, Adler M, Pearson A, Gelman F, Bar-Or I, John SG, Eckert W (2011) Geochemical evidence for iron-mediated anaerobic oxidation of methane. Limnology and Oceanography, 56, 1536–1544.

Sivan O, Antler G, Turchyn AV, Marlow JJ, Orphan VJ (2014) Iron oxides stimulate sulfate-driven anaerobic methane oxidation in seeps. Proceedings of the National Academy of Sciences, 111, 4139–4147.

Stolz JF, Ellis DJ, Blum JS, Ahmann D, Lovley DR, Oremland RS (1999) Note: Sulfurospirillum barnesii sp. nov. and Sulfurospirillum arsenophilum sp. nov., new members of the Sulfurospirillum clade of the ε-Proteobacteria. International Journal of Systematic and Evolutionary Microbiology, 49, 1177–1180.

Stookey LL (1970) Ferrozine–-a new spectrophotometric reagent for iron. Analytical Chemistry, 42, 779–781.

Stüeken EE, Buick R, Anbar AD (2015) Selenium isotopes support free O2 in the latest Archean. Geology, 43, 259–262.

Sturm A, Fowle DA, Jones C, Leslie KL, Nomosatryo S, Henny C, Canfield DE, Crowe S (2015) Rates and pathways of methane oxidation in ferruginous Lake Matano, Indonesia. Biogeosciences Discussions, 12, 1–34.

Ueno Y, Yamada K, Yoshida N, Maruyama S, Isozaki Y (2006) Evidence from fluid inclusions for microbial methanogenesis in the early Archaean era. Nature, 440, 516–519.

Vargas M, Kashefi K, Blunt-Harris EL, Lovley DR (1998) Microbiological evidence for Fe(III) reduction on early Earth. Nature, 395, 65–67.

Willems A (2014) The family Comamonadaceae. In: The Prokaryotes. Springer, pp 777–851.

Zavarzina DG, Sokolova TG, Tourova TP, Chernyh NA, Kostrikina NA, Bonch-Osmolovskaya EA (2007) Thermincola ferriacetica sp. nov., a new anaerobic, thermophilic, facultatively chemolithoautotrophic bacterium capable of dissimilatory Fe(III) reduction. Extremophiles, 11, 1–7.

Zegeye A, Bonneville S, Benning LG, Sturm A, Fowle DA, Jones C, Canfield DE, Ruby C, Maclean LC, Nomosatryo S (2012) Green rust formation controls nutrient availability in a ferruginous water column. Geology, 40, 599–602.

Zehnder AJ, Brock TD (1979) Methane formation and methane oxidation by methanogenic bacteria. Journal of Bacteriology, 137, 420–432.

Zehnder AJ, Brock TD (1980) Anaerobic methane oxidation: occurrence and ecology. Applied and Environmental Microbiology, 39, 194–204.

Zhou S, Xu J, Yang G, Zhuang L (2014) Methanogenesis affected by the co-occurrence of iron (III) oxides and humic substances. FEMS Microbiology Ecology, 88, 107–120.

